# Asymmetric neural entrainment at resonance frequencies underlies unilateral spatial neglect

**DOI:** 10.1101/2024.10.22.619765

**Authors:** Yuka O. Okazaki, Noriaki Hattori, Teiji Kawano, Megumi Hatakenaka, Ichiro Miyai, Keiichi Kitajo

**Affiliations:** Division of Neural Dynamics, Department of System Neuroscience, National Institute for Physiological Sciences, National Institutes of Natural Sciences, Okazaki, Japan; Physiological Sciences Program, Department of Advanced Studies, Graduate Institute for Advanced Studies, Graduate University for Advanced Studies (SOKENDAI), Okazaki, Japan; Department of Rehabilitation, University of Toyama, Toyama, Japan; Neurorehabilitation Research Institute, Morinomiya Hospital, Osaka, Japan; RIKEN CBS-TOYOTA Collaboration Center, RIKEN Center for Brain Science, Wako, Japan

**Keywords:** Alpha oscillations, Hemispheric imbalance, Steady-state visual evoked potentials

## Abstract

Unilateral spatial neglect (USN) is a common consequence of right-hemisphere stroke, traditionally attributed to structural lesions and dysfunctional attention networks. However, the brain is fundamentally a rhythmic and dynamical system, and how disrupted neural synchronization underlies USN remains unknown. We recorded steady-state visual evoked potentials (SSVEPs; 3 - 30 Hz flicker) in stroke patients with USN, without USN, and healthy controls. Only the USN group exhibited significant hemispheric asymmetry at 9 Hz, driven by exaggerated responses in the intact hemisphere rather than suppression in the lesioned hemisphere. This effect appeared only during stimulation, not at rest, indicating its specificity to sensory processing. The enhanced 9 Hz entrainment in the intact hemisphere was accompanied by increased phase-amplitude coupling (PAC) between alpha phase and gamma amplitude, reflecting systematic coordination of high-frequency activity. Transfer entropy analysis further revealed increased feedforward information flow from the right visual to the left frontal cortex, highlighting large-scale asymmetry. To explore the mechanism underlying this frequency-specific bias, we implemented a coupled-oscillator model. The model showed that the hemispheric asymmetry arises from resonance between intrinsic alpha rhythms and external input, amplified by asymmetric right-to-left interhemispheric coupling. These findings suggest that USN arises from a selective impairment of alpha-band synchrony capacity. This study offers a novel framework conceptualizing USN as a disorder of disrupted oscillatory dynamics underlying spatial attention, and points toward frequency-specific neuromodulatory intervention as a potential therapeutic approach.

**Significant statement:** The brain behaves as a metronome-like oscillator ensemble. When a rhythmic flicker—a cortical “tuning fork” matching the visual system’s intrinsic alpha frequency—is applied, it serves as a probe that visualizes the brain’s internal state. Under normal conditions, neuronal oscillations in both hemispheres are moderately entrained at this frequency. In unilateral spatial neglect (USN), this resonance displays a distinctive asymmetry: the intact hemisphere becomes over-amplified, whereas the lesioned hemisphere shows no enhancement. A coupled-oscillator model shows that asymmetric interhemispheric coupling selectively boosts this intrinsic alpha resonance unilaterally. This biased resonance reflects a destabilization of the dynamic coordination of attention networks, offering a novel mechanistic view of spatial neglect in USN and pointing toward frequency-specific neuromodulation as a promising therapeutic strategy.

## Introduction

Left unilateral spatial neglect (USN) is a neurological disorder in which patients fail to perceive stimuli on the left side of space despite an intact visual field. This condition is associated with damage to the right dorsal attention network, particularly the right inferior parietal lobule (Corbetta et al. 2005; Mort et al. 2003), and the superior temporal gyrus (Karnath 2001: 21; Karnath, Ferber, and Himmelbach 2001), as well as disconnection in occipital and frontal white matter pathways (Thiebaut de Schotten et al. 2005). The dorsal attention network (Corbetta and Shulman 2002), which includes the frontal eye fields and bilateral parietal sulci, encodes spatial maps of the contralateral visual field in each hemisphere. These maps are typically balanced through reciprocal inhibitory interactions(Grefkes and Fink 2011). Disruption of this network leads to hyperactivity of the structurally intact left hemispheric circuits, which influences bottom-up sensory responses and causes a rightward bias in perception (Corbetta et al. 2005; He et al. 2007).

Traditionally, USN has been understood in terms of structural or static disconnection. However, the brain is fundamentally a dynamical and frequency-organized system, and neural oscillations, generated by the synchronized activity of neuronal populations, are a key mechanism for temporally coordinating information flow across brain regions. Indeed, neural oscillations flexibly regulate sensory and attentional processing by dynamically adjusting intrinsic and stimulus-driven phase relationships (Busch, Dubois, and VanRullen 2009; Mathewson et al. 2009; Lakatos et al. 2013; Lakatos et al. 2008; Schroeder and Lakatos 2009; Fries 2015). From this perspective, USN may not only involve anatomical disconnection but also an imbalance in the hemispheric capacity for oscillatory synchronization. Yet, despite the critical role of rhythmic synchronization in brain function, such frequency-specific asymmetries have not been systematically investigated in USN.

To address this, we employed steady-state evoked responses (SSERs) to visual stimuli as a probe to visualize the brain’s frequency-dependent synchronization capacity. SSERs are elicited by rhythmic sensory stimulation and provide a measure of stimulus-locked neural responses. When the stimulation frequency is close to the system’s intrinsic resonance frequency, these responses may include an entrainment component, reflecting the alignment of endogenous oscillations to external input (Pikovsky, Rosenblum, and Kurths 2003; Okazaki et al. 2021). Moreover, prior studies have shown frequency-specific reductions in SSERs, both visual and auditory, in individuals with schizophrenia (Krishnan et al. 2005; Kwon et al. 1999) and autism (Wilson et al. 2007; Rojas et al. 2011). Consequently, SSER abnormalities may reflect impairments in underlying oscillatory mechanisms.

In this study, we presented 3–30 Hz flickering stimuli and recorded steady-state visual evoked potentials (SSVEPs) in stroke patients with and without USN, as well as in healthy controls. None of the patients exhibited clinically evident hemianopsia, and early-stage visual responses in the lesioned hemisphere were not expected to be significantly impaired. These factors minimized the possibility that observed differences could stem from low-level visual deficits. We hypothesized that frequency-specific stimulus-locked synchronization dynamics in response to rhythmic stimulation would be selectively disrupted in one hemisphere, resulting in an interhemispheric imbalance in USN. Furthermore, motivated by the empirical asymmetry, we developed a computational model of coupled neural oscillators (Kuramoto 1984; Sakaguchi 1988) to test which factors could account for the emergence of frequency-specific differences in oscillatory responses.

## Materials and Methods

### Participants

The study consisted of three groups: USN patients (n = 21), stroke patients without USN (n = 38), and healthy control participants (n = 17). We recruited patients who had been admitted to the Kaifukuki (convalescent) Rehabilitation Ward of Morinomiya Hospital. The inclusion criteria for stroke patients were: (1) male, (2) first-episode stroke, and (3) right hemisphere cortical/subcortical infarction (lesion overlap maps in Supplementary Figure 1), without clinically evident hemianopsia evaluated by the confrontation method. While lesions partially extended to the posterior region in two patients, they exhibited preserved neurophysiological responses to light stimulation, and were therefore included in the analysis. For healthy participants, the inclusion criteria were: (1) male to match the patient group, (2) no history of stroke, and (3) full scores on all neuropsychological assessments. All participants provided written informed consent. The study was approved by the Institutional Review Boards of RIKEN, the National Institutes of Natural Sciences, and Morinomiya Hospital.

### Clinical assessment of stroke

Electroencephalography (EEG) measurements and clinical assessments were performed by trained investigators approximately 1 month after stroke onset. The diagnosis of USN was comprehensively determined by the attending physician based on regular clinical examinations, including the Behavioral Inattention Test (BIT) (Wilson, Cockburn, and Halligan 1987). Motor impairment was assessed using the Fugl-Meyer Assessment upper extremity (FMA-UE) and lower extremity (FMA-LE) (Fugl-Meyer et al. 1975), and activities of daily living were assessed using the Functional Independence Measure motor (FIM-M) and cognitive (FIM-C) scores (Keith et al. 1987). All clinical assessments were performed by trained healthcare professionals before EEG analysis, so the evaluators were blind to the EEG analysis results. For all scales, a lower score indicates greater disability. To match the severity of the USN and non-USN groups, we selected 21 non-USN patients in ascending order of the Mahalanobis square distance, *d_i_^2^*, to the USN patient within a four-dimensional space of the clinical score:

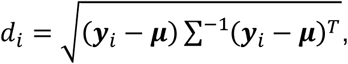

where ***y****_i_* is the *i*-th non-USN patient’s clinical scores and ***μ*** and ***Σ*** are the mean and covariance matrix of the USN samples, respectively. The demographic information and clinical scores of each group are listed in Table 1. A Mann–Whitney U test confirmed that there were no significant differences between the groups in any of the scores.

**Table 1.** Demographic information of each group and clinical scores of the USN and non-USN patients matched by Mahalanobis distance.

| Variables | Median (Q1–Q3) |  |  | *p*-value<br>(*U*-test) |
| --- | --- | --- | --- | --- |
|  | Non-USN | USN | Healthy control |  |
| Number of participants | 21 | 21 (16) | 17 | — |
| Sex | Male | Male | Male | — |
| Age (years) | 69.0<br>(62.8–78.2) | 67.0<br>(54.8–77.5) | 67.0<br>(58.072.2) | 0.21, 0.91<br>(USN vs. non-USN, USN vs. Healthy) |
| Cortical/subcortical lesion side | Right | Right | — | — |
| EEG recording (days poststroke) | 32.0<br>(21.5–41.0) | 38.0<br>(32.5–43.0) | — | 0.19 |
| FMA-UE (0–66) | 49.0<br>(34.2–61.2) | 53.0<br>(12.8–62.2) | — | 0.93 |
| FMA-LE (0–34) | 26.0<br>(22.5–30.0) | 28.0<br>(16.2–31.0) | — | 0.92 |
| FIM-M (13–91) | 64.0<br>(45.0–71.5) | 47.0<br>(32.8–72.8) | — | 0.23 |
| FIM-C (5–35) | 30.0<br>(26.8–31.2) | 29.0<br>(22.0–30.0) | — | 0.13 |
| BIT (0–222) | — | 204.5<br>(195.5–216.5) | — | — |

### Experimental Design

EEG was recorded from 19 Ag/AgCl electrodes in the 10-20 system configuration using a NeuroFax EEG 1224 system (Nihon Kohden, Tokyo, Japan; sampling rate: 500 Hz; bandpass filter: 0.53–120 Hz; ground electrode: forehead; reference: left earlobe). Visual stimulation was delivered under an eyes-closed resting condition to minimize blink artifacts and reduce direct exposure to the high-intensity strobe light. A strobe light flickered at frequencies ranging from 3 to 30 Hz in 3-Hz steps. Each frequency flicker stimulus was presented for 10 s with a 10-s interval (Figure 1).

**Figure 1.**
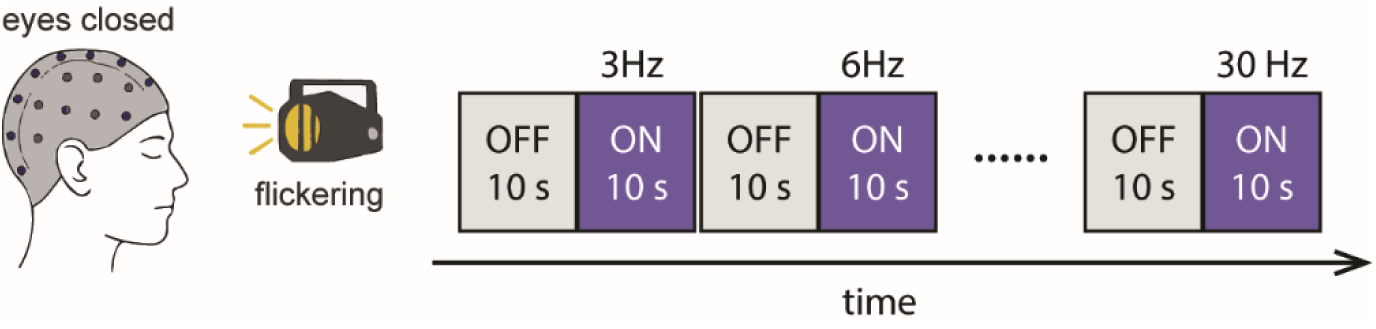
Experimental paradigm. The strobe light flickered for 10 seconds in 3-Hz steps, from 3 Hz to 30 Hz, with 10-second inter-train intervals. Participants were lying down with their eyes closed during the EEG recording.

### Preprocessing

All EEG analyses were implemented using custom scripts in MATLAB (MathWorks, Natick, MA, USA) and the FieldTrip toolbox (Oostenveld et al. 2011). The EEG signals were first re-referenced offline to the average of the right and left earlobe signals. We applied wavelet-enhanced independent component analysis (wICA) to eliminate intermittent high-amplitude artifacts (Castellanos and Makarov 2006). This method decomposes the EEG signals into independent components (ICs), and a discrete wavelet transform is applied to each IC. Wavelet coefficients defined as artifacts based on the level-dependent threshold of the wavelet decomposition are zeroed, and artifact-free ICs are extracted from the inverse wavelet transform on this result. Thus, the entire IC, which contains a significant amount of neural signals, need not be excluded, as in the case of conventional ICA denoising.

### Steady-state visual evoked potential (SSVEP)

To evaluate the neural response during the flickering stimulation, the frequency power spectrum for each stimulus condition was obtained using the Fourier transform (Hanning window, frequency range: 1–35 Hz). The Fourier transform was applied to the data with a 1-s window during the 10-s flicker interval and averaged for each stimulus frequency condition. For the resting-state interval, power was estimated using the same procedure from the ‘off’ interval immediately preceding the 3-Hz flicker block (see Fig. 1). SSVEP power at the fundamental (stimulated) frequency was then extracted for subsequent analyses. Because the fundamental SSVEP component is typically maximal over occipital electrodes in healthy participants (Norcia et al. 2015), we evaluated occipital electrodes (O1/O2) as primary sites. The scalp topography of the mean fundamental-frequency SSVEP power averaged across stimulation conditions is shown in Supplementary Fig. 2.

### Phase-amplitude coupling (PAC)

We evaluated PAC using the modulation index (MI) (Tort et al. 2010) to determine how high-frequency oscillations are co-modulated by a low-frequency phase when neural oscillations are entrained to rhythmic external stimuli. Instantaneous phase and amplitude were extracted via wavelet transform at the center frequency, *f*, and time, *t*, with the standard deviation (SD), *σ_f_ = 4f/m*, *σ_t_ = m/2*π*f*, and constant, *m,* set to 4 (Lachaux et al. 2000). We focused on the PAC between the alpha (6–12 Hz) phase and the gamma (30–55 Hz) amplitude for 10 s during the 9-Hz flickering stimulus.

To derive the MI for each frequency pair of α phase and γ amplitude, the given α frequency phase was divided into 20 bins, and the mean γ amplitude in each bin was calculated. This provided a phase-amplitude distribution for each frequency pair, and the MI was obtained according to the Kullback-Leibler divergence between the observed distribution and the circular uniform distribution. An MI of 0 indicates that the amplitudes are the same for all phase bins (i.e., a circular uniform distribution) and that there is zero PAC, whereas an MI of 1 indicates maximum coupling.

### Transfer entropy (TE)

To quantitatively investigate the direction and strength of information transfer between brain regions in response to flickering stimuli, we performed TE analysis (Schreiber 2000). After applying an 8.5–9.5 Hz FIR bandpass filter to the EEG responses to 9-Hz flickering stimuli, we extracted the instantaneous amplitude (amplitude envelope) from the analytic signal using the Hilbert transform. Because this amplitude envelope fluctuates nonperiodically at a slower timescale than the carrier 9-Hz oscillation, it provides a broadband measure of signal dynamics while minimizing the strong autocorrelation inherent in narrow-band periodic signals that can spuriously inflate TE estimates (Daube, Gross, and Ince 2022). TE was then calculated for all pairs of electrodes for these amplitude data during the flickering stimulus interval. To estimate the probability distributions required for TE calculation, we generated histograms for the amplitude data, with bin sizes according to Sturges’ rule (Legg et al. 2013). *T_Y→X_* quantifies the extent to which knowledge of the state of system *Y* at time *t* contributes to predicting the one-step future state of system *X* at time *t+τ*, beyond the predictability provided by the state of *X* at time *t* alone. Importantly, because TE evaluates time-lagged prediction (from Y(t) to X(t+τ)), it is not designed to capture zero-lag common-source correlations (i.e., volume conduction) and therefore characterizes directed, nonzero-lag dependencies rather than instantaneous coupling. The formula for TE is given by:

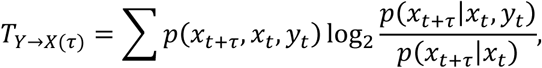

where *p*(*x_t_*_+*τ*_, *x_t_*, *y_t_*) denotes the joint probability of observing the future state of *X* at time *t+τ* along with the states of *X* and *Y* at time *t.* Here, *x* and *y* represent the observed data derived from the systems *X* and *Y*, respectively. The terms *p*(*x_t_*_+*τ*_|*x_t_*, *y_t_*) and *p*(*x_t_*_+*τ*_|*x_t_*) denote the conditional probabilities of the one-step future state of *X,* given its state at time *t* with and without considering the state of *Y* at time *t*.

In this analysis, amplitude data from the first 0.5 s of the flickering stimulus interval were used as past observations, *x_t_*, *y_t_*, and amplitude data from one sample ahead in time were used as future observations, *x_t+τ_*. TE was calculated by varying the delay, *τ*, from 0 to 5 s in 0.1-s steps. The optimal delay *τ* was determined as the value that maximized *T*(*τ*).

### Statistical analysis

The SSVEP power at the fundamental (stimulated) frequency *f* was analyzed for each condition as the dependent variable using a linear mixed model in SPSS, with hemisphere (left and right), flicker frequency (3, 6, 9, 12, 15, 18, 21, 24, 27, and 30 Hz), and group (USN patients, non-USN patients, and healthy controls) as fixed factors and subjects as random factors. Outliers exceeding 1.5 times over the interquartile range were excluded.

To statistically test for differences in PAC between the left and right hemispheres, we performed a cluster-based permutation test (Maris and Oostenveld 2007) using Fieldtrip. The MI was calculated for each permutation dataset generated by random reassignment of O1 and O2 electrode labels. For each permutation, clusters were identified from adjacent data points showing statistically significant hemispheric differences in MI, and the sum of the *t*-values within these clusters was determined as a cluster-level statistic. Only the largest value of the cluster-level statistic obtained for each permutation was selected, and this procedure was repeated 1000 times to construct the distribution of the cluster-level statistic under the null hypothesis. A cluster of hemispheric differences in the original data was considered statistically significant if it exceeded the 95th percentile of this distribution.

### The coupled phase oscillator model

To investigate potential mechanisms underlying our experimental findings, we employed the Kuramoto model (Kuramoto 1984) as a theoretical framework for neural synchronization. The model consists of bilateral coupled phase oscillators and an external periodic force (Sakaguchi 1988). Two coupled populations of oscillators (*N* = 50 each) are represented by phase vectors ***Φ*_L_**= (*ϕ*_L,1_, …, *ϕ*_L,n_) and ***Φ*_R_** = (*ϕ*_R,1_, …, *ϕ*_R,n_) for the left and right hemispheres, respectively. The dynamics of each oscillator are given by:

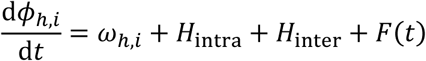

The natural frequencies *ω_h,i_* follow a Gaussian distribution with a mean of 9 Hz and a standard deviation of 2 Hz. In one configuration, we set the mean natural frequency of the right hemisphere to 9 Hz to examine the effect of altered intrinsic frequency in the affected hemisphere. The intrahemispheric coupling *H*_intra_ and interhemispheric coupling *H*_inter_ terms are defined as:

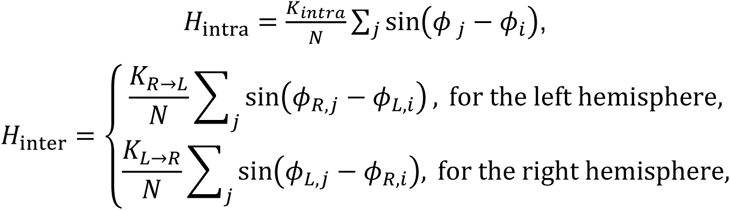

where *ϕ_,i_* represents the phase of the *i*-th oscillator in left and right hemispheres. *K*_intra_ represents intrahemispheric coupling strength, and *K_R→L_* and *K_L→R_* represent interhemispheric coupling strengths from right to left and left to right hemispheres, respectively. The external forcing term is given by:

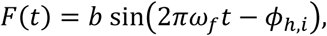

where amplitude and frequency of the external periodic force are set to *b* = 2 and *ω_f_ ∈* {*3, 6, …, 30*} Hz. To quantify the degree of synchronization, we computed the instantaneous order parameter within each hemisphere:

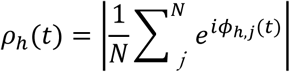

We performed 21 independent simulations, each corresponding to a virtual subject with different initial phases randomly drawn from a uniform distribution [0, 2pi] and natural frequencies sampled from Gaussian distributions. The order parameter was averaged over 10 seconds with continuous stimulation. The numerical simulation used the 4th-5th-order Runge-Kutta method with adaptive time steps, as implemented by MATLAB’s ode45 function.

## Results

### SSVEP power in the affected hemisphere

To confirm the visual cortex of the affected (right) hemisphere could respond to visual stimuli, we plotted SSVEP power at the O2 electrode as a function of the flicker and response frequencies (Figure 2). When compared with the power spectrum of the baseline period, a clear fundamental can be seen on the diagonal (black dashed line), which indicates that the visual cortex of the affected hemisphere was precisely enhanced at the flicker frequency.

**Figure 2.**
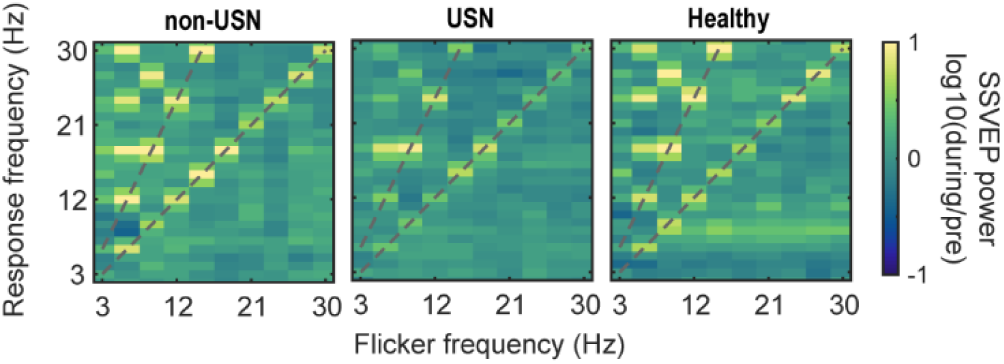
SSVEP power for different responses and flicker frequencies. The power spectrum at the response frequency (y-axis) is shown for the O2 electrode (affected hemisphere in patients) as a function of flicker frequency (x-axis). Spectra are normalized to the baseline from the resting-state interval. The fundamental frequency and the first harmonic (gray dashed lines) are evident across groups.

### Hemispheric imbalance of the SSVEP

The SSVEP power of the affected (right) hemisphere (O2 electrode) was compared with that of the intact (left) hemisphere (O1 electrode) to investigate hemispheric balance in frequency responses. We found a significant hemisphere × flicker frequency × group interaction (*F* = 2.09, *p* < 0.005). To further investigate this interaction, a simple main effects test was conducted for hemisphere differences at each frequency within each group, with a Bonferroni correction applied for multiple comparisons. Within the USN group, SSVEP power of the intact hemisphere was significantly greater than that of the affected hemisphere at 9 Hz (*F* = 62.36, *p* < 0.001). No significant hemisphere differences were observed at other frequencies within the USN group. In the non-USN and healthy control groups, there were no significant hemisphere differences at any frequency after Bonferroni correction. Figure 3 shows the hemispheric difference in SSVEP power (O1 - O2) corresponding to stimulation frequency. This suggests that frequency-specific hemispheric bias is the factor that separates USN from non-USN patients.

**Figure 3.**
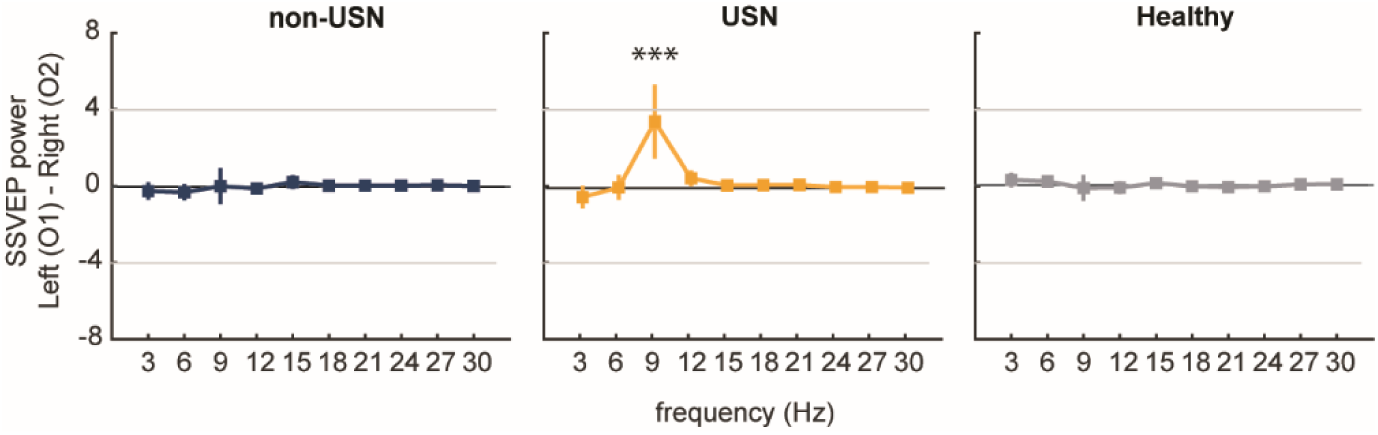
Hemispheric differences in SSVEP power at different flicker frequencies. Power in the intact hemisphere was stronger than that in the affected hemisphere with 9-Hz stimulation (\*\*\**p* < 0.001) in the USN group only (middle panel). Error bars represent 2 standard errors.

We applied the same analysis for the central (C3, C4) and frontal (F3, F4) electrodes to verify the frequency response of hemispheric bias outside the visual area. Figure 4 shows the difference in SSVEPs between C3 and C4 electrodes (Figure 4A) and between F3 and F4 electrodes (Figure 4B) at each frequency. We found no significant main effects for hemispheres or interactions in either the central or frontal regions.

**Figure 4.**
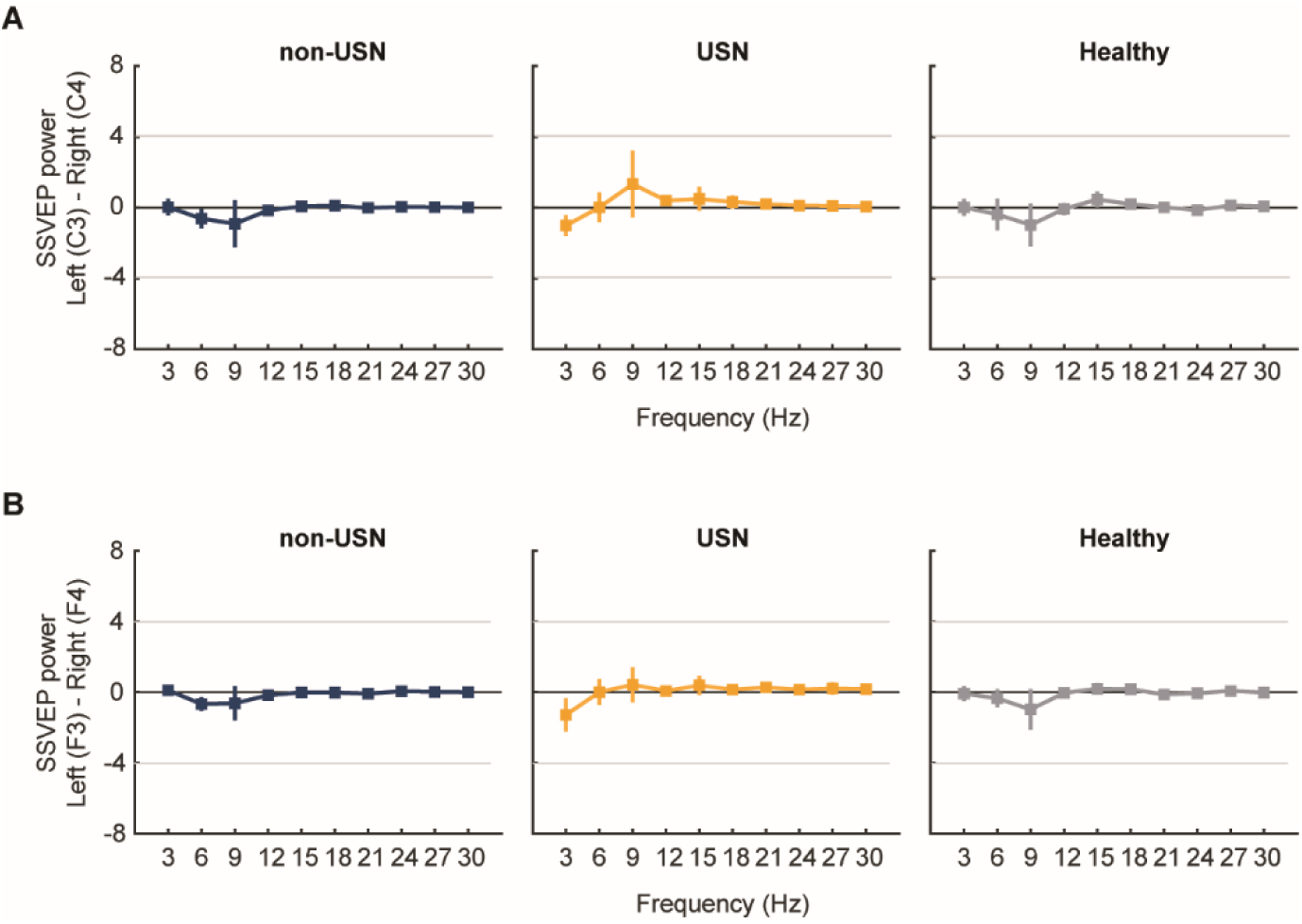
Hemispheric differences in SSVEP power at different flicker frequencies. **(A)** No hemispheric difference between C3 and C4 electrodes. **(B)** No hemispheric difference between F3 and F4 electrodes. Error bars represent 2 standard errors.

### Hyperactivity of the intact hemisphere

To determine whether the hemispheric imbalance of SSVEP power confined to 9 Hz in the USN group was attributed to the intact (left) or affected (right) hemisphere, we compared SSVEP power at 9 Hz in the corresponding hemisphere in the healthy controls (Figure 5A). A simple main effects test showed a significant effect of group on SSVEP power in the left hemisphere (*F* = 6.66, *p* < 0.005) and in the right hemisphere (*F* = 10.58, *p* < 0.001). Post-hoc comparisons with Bonferroni correction indicated that in the left hemisphere, both the USN and the non-USN group had significantly higher SSVEP power than the healthy control group (p < 0.005 and *p* < 0.05, respectively). In the right hemisphere, the non-USN group showed significantly higher SSVEP power than the healthy control group (*p* < 0.005), while the USN group did not differ significantly from the healthy controls. We also examined the resting state interval (i.e., before the photo flicker session) and found no significant differences in either the USN or non-USN group (Figure 5B). Taken together, there was no major abnormality in the spontaneous generation of alpha oscillations in the USN group; rather, it was the capacity to synchronize these oscillations with other external oscillations that differed between the hemispheres in the USN group.

**Figure 5.**
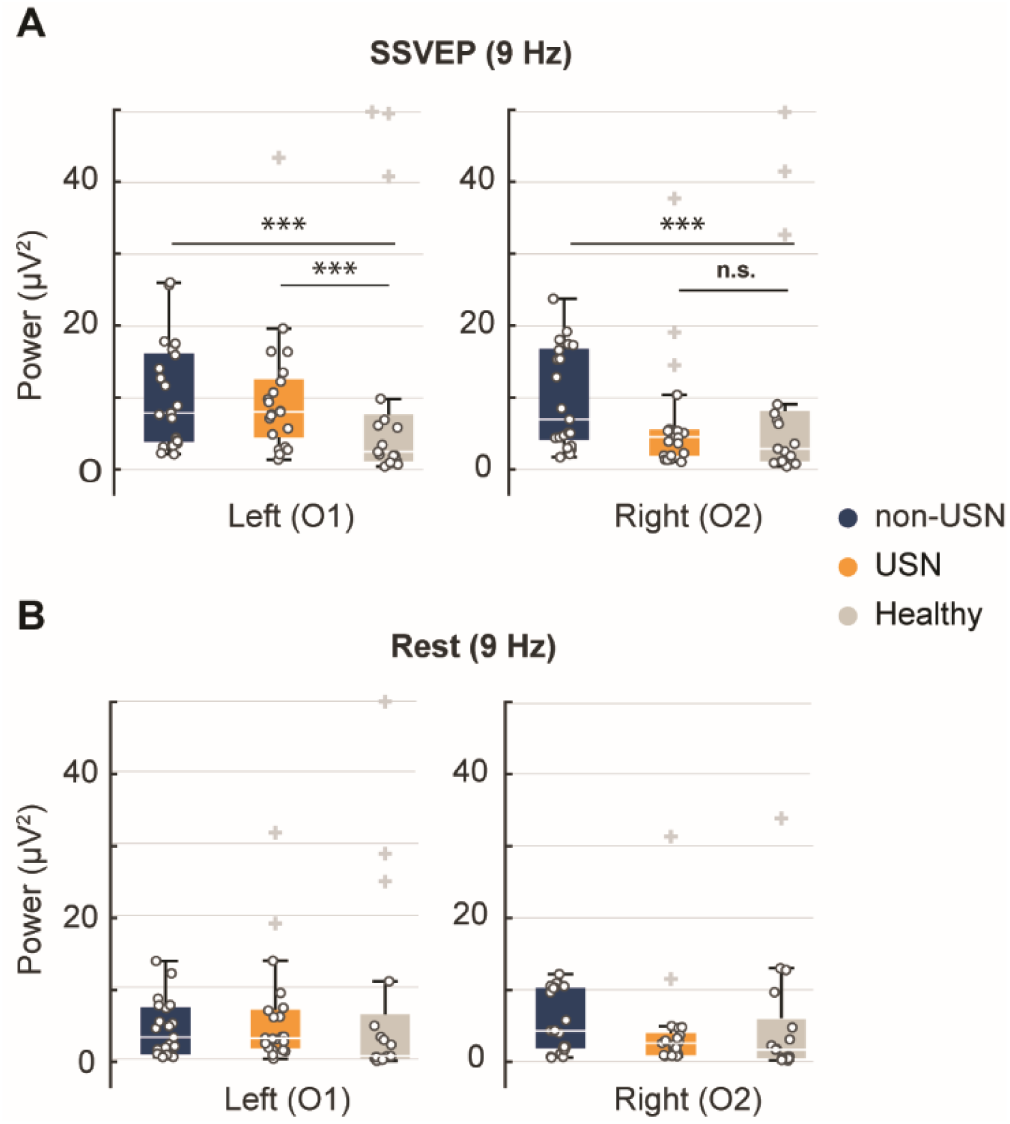
SSVEP power during stimulation at 9 Hz and rest. **(A)** Box plots indicate the power of SSVEP responses in the intact (O1 electrode) and affected (O2 electrode) hemispheres. A significantly higher response power was observed in the intact hemisphere of the USN group than in the left hemisphere of the healthy control group. The non-USN patients also displayed higher alpha power in both hemispheres than healthy controls without hemispheric bias (***p < 0.001). **(B)** During the resting state interval, no significant differences in alpha power were observed between groups. Box plots depict medians and 25th/75th percentiles, with whiskers showing the 10th/90th percentiles overlaid onto individual data points (circles). The + represents outliers. Outliers exceeding a power value of 50 were truncated and plotted at the 50 mark.

### Hemispheric imbalance of PAC

We investigated PAC given that higher frequency oscillations may be cooperatively modulated when neural oscillations are entrained by rhythmic external stimuli. PAC was assessed using the MI for 9-Hz flickering stimuli (Figure 6). We hypothesized that the intact left hemisphere, which showed enhanced SSVEP, would exhibit stronger PAC. Thus, we performed a cluster-based permutation test to examine hemispheric differences. Results showed a significant hemispheric bias in PAC between the alpha (7–12 Hz) phase and gamma (35–45 Hz) amplitude in the USN group. Specifically, the MI was higher in the intact hemisphere than in the affected hemisphere (*p* < 0.05, one-tailed) peaking at the 9-Hz phase. These effects were not observed in the non-USN or healthy control group. This suggests that gamma activity is modulated by the alpha phase aligned to the 9-Hz flicker stimulation and that this modulation is particularly enhanced in the intact hemisphere of the USN group.

**Figure 6.**
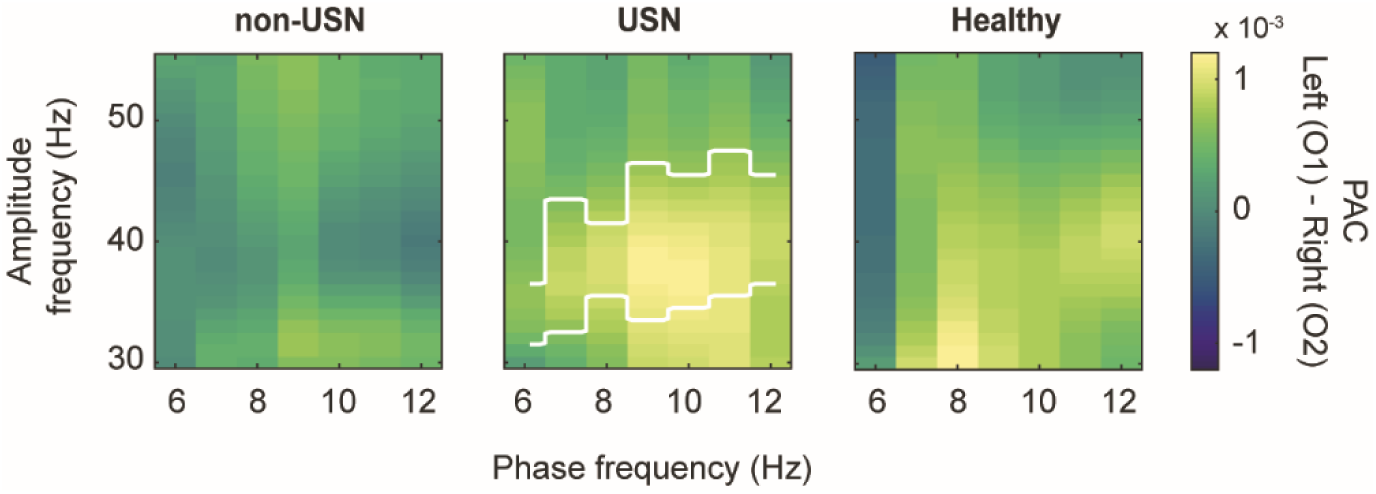
Hemispheric differences in PAC in response to 9-Hz flickering stimuli. The MI measuring the coupling between the alpha phase (7–12 Hz) and gamma power (35–45 Hz) represents the hemispheric asymmetry in the USN group (indicated by the black outline). No such pattern was present in the non-USN or healthy control group.

### Hemispheric imbalance of TE

In USN and non-USN patients with right hemisphere damage, we aimed to clarify how information is transferred from the responsive visual cortex to other brain regions. Here, we used TE to examine the pattern of information transfer from the O1 and O2 electrodes to the left and right frontal electrodes (Figure 7). Results showed that in the USN group, there was significantly greater information flow from the right visual area (O2 electrode) than from the left visual area (O1 electrode) to the intact left frontal areas (*t*(20) = −2.7, *p* < 0.05, paired *t*-test). This enhancement was significantly greater than that in the healthy control group (*t*(36) = −2.89, *p* < 0.01, unpaired *t*-test; Figure 7A, left panel). This indicates a hemispheric bias in information transfer in the feedforward direction in USN patients. In contrast, no such bias was observed in non-USN patients (*t*(20) = −0.47, *p* = 0.64, paired *t*-test) or healthy controls (*t*(16) = 1.42, *p* = 0.174, paired *t*-test). There were no significant differences in information flow from the O1/O2 electrodes to the affected right frontal areas (Figure 7A, right panel) or in the feedback direction (i.e., from the left/right frontal areas to the O1/O2 electrodes) in any group (Figure 7B). These findings suggest that the distinct clinical symptoms of USN are not solely due to biased local brain activity but are closely related to asymmetric information transfer, especially in the feedforward direction.

**Figure 7.**
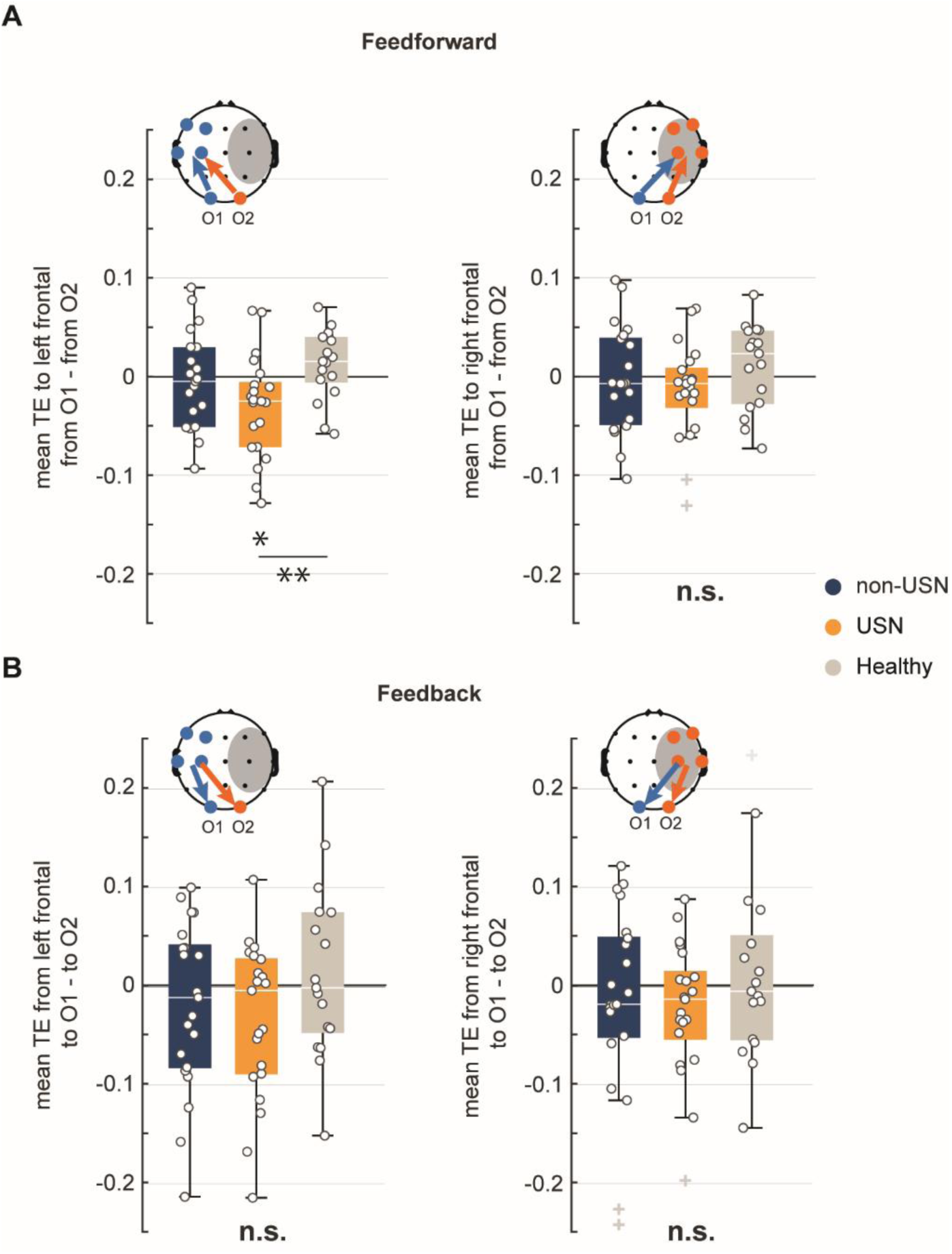
Hemispheric differences in information flow between the visual and frontal areas in response to 9-Hz flickering stimuli. **(A)** Box plots show left-right hemispheric asymmetry in transfer entropy (TE) from visual to frontal areas (feedforward direction). In the USN group, information flow from the right visual cortex (O2) to the left frontal area (left panel) was significantly stronger than from the left visual cortex (O1) to the left frontal area (\**p* < 0.05), and this asymmetry was significantly greater than in healthy controls (\*\**p* < 0.01). No significant hemispheric differences were observed in information flow to the right frontal area (right panel). **(B)** Box plots show left-right hemispheric asymmetry in TE from frontal to visual areas (feedback direction), with no significant asymmetry detected in any group for either left or right frontal areas. Arrows on topographical plots indicate the direction of information flow. Box plots show medians and 25th/75th percentiles, with whiskers at 10th/90th percentiles. The + represents outliers.

### Correlation between SSVEP power and USN severity

To investigate the relationship between neural synchronization capacity and symptom severity in USN patients, we performed Pearson’s correlation analysis between SSVEP power and BIT scores. Results showed a significant positive correlation between SSVEP power at 9 Hz in the right hemisphere (O2 electrode) and BIT score (Pearson’s *r* = 0.70, *p* < 0.01). That is, the higher the SSVEP power at 9 Hz in the right hemisphere, the milder the USN symptoms. In contrast, there was no significant correlation in the left hemisphere (O1 electrode; Pearson’s *r* = 0.38, *p* = 0.18; Figure 8A). Additionally, to examine the frequency-specific nature of the relationship between SSVEP power and USN severity, we analyzed correlations across all stimulation frequencies. Significant correlations between SSVEP power and BIT score were observed only at 9 and 12 Hz in the right hemisphere (Pearson’s *r* = 0.70, *p* < 0.01, for both frequencies, FDR-corrected; Figure 8B). No significant correlations were found at any frequency in the left hemisphere (O1 electrode). Our results showed that USN patients maintained 9 Hz SSVEP power in the right hemisphere comparable to healthy controls. However, within the USN group, this right hemisphere activity significantly correlated with symptom severity, with higher SSVEP power associated with milder USN symptoms. In addition, correlations between BIT and the hemispheric imbalance of SSVEP power are provided in the Supplementary Information (Supplementary Fig. 3A). We also report, as additional exploratory analyses, correlations between BIT and hemispheric imbalance measures derived from PAC and TE (Supplementary Fig. 3B and 3C). None of these correlations reached statistical significance after correction, although the feedforward TE imbalance to the left frontal region showed a marginal uncorrected association with BIT score (*r* = -0.51, uncorrected *p* = 0.05).

**Figure 8.**
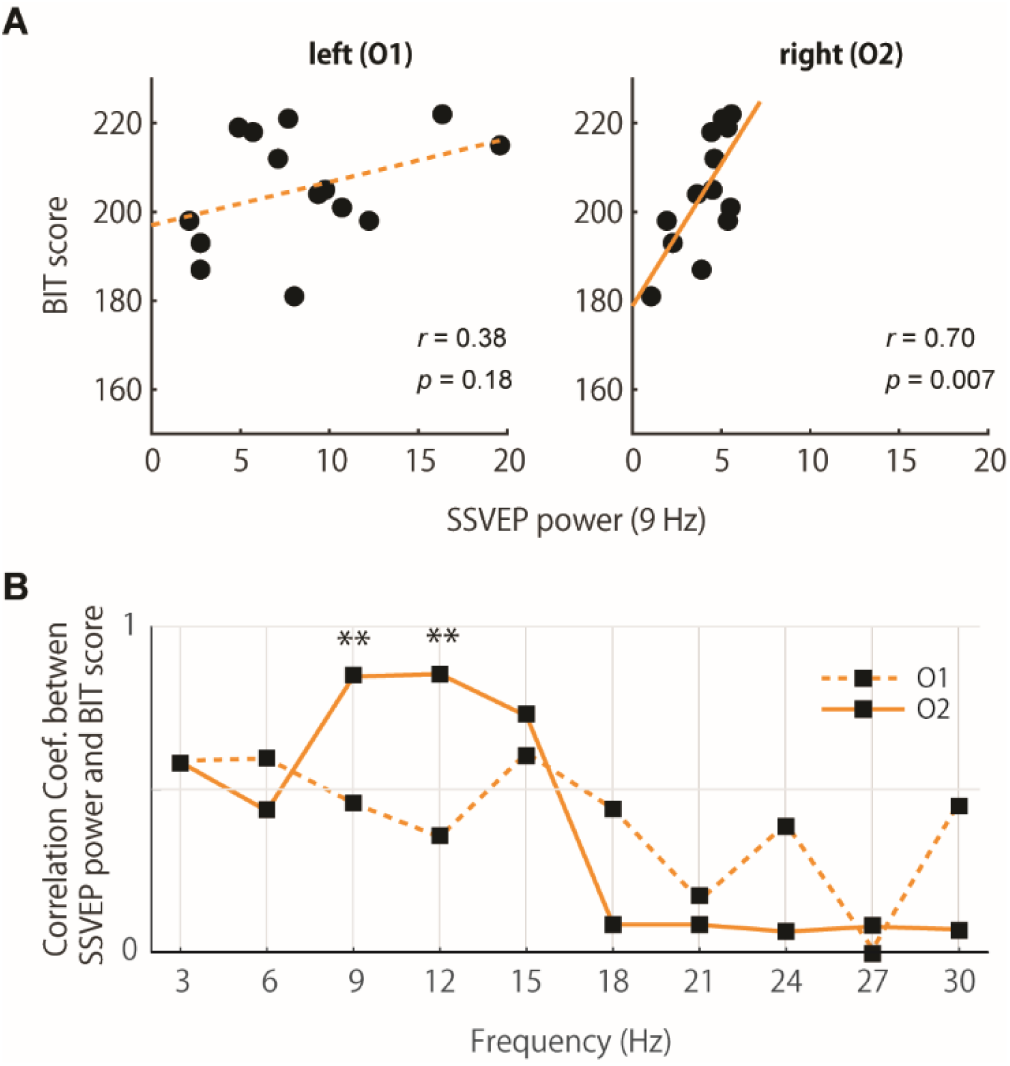
Correlation between SSVEP power and BIT score in USN patients. **(A)** Scatter plots represent the relationship between SSVEP power at 9 Hz and BIT score for the left hemisphere (O1 electrode, left panel) and right hemisphere (O2 electrode, right panel). A significant positive correlation was found in the right hemisphere (*r* = 0.70, *p* < 0.007) but not in the left hemisphere (*r* = 0.38, *p* = 0.18). **(B)** Correlation coefficients between SSVEP power and BIT score across frequencies (3–30 Hz) for each hemisphere. Significant correlations were observed in the right hemisphere only at 9 and 12 Hz after FDR correction (\*\**p* < 0.01).

### Hemispheric Imbalance in Model Simulations

To explain the observed hemispheric imbalance in USN patients (Figure 3, middle), we explored three potential mechanisms using a computational model. From a base configuration (*ω_L_= ω_R_=* 9±2 Hz, *K_L_ = K_R_ = K_L→R_ = K_R→L_* = 0.2), we modified specific parameters. In the first configuration, we introduced asymmetric interhemispheric coupling (*K_R→L_* = 0.8). This configuration most closely reproduced our experimental findings, showing a significant hemispheric bias specifically at 9 Hz (p < 0.05, FDR corrected), while maintaining balanced responses at other frequencies (Figure 9A). In the second configuration, we increased intrahemispheric coupling strength in the intact hemisphere (*K_L_* = 0.8) while maintaining the baseline coupling in the affected hemisphere (*K_R_* = 0.2), based on our experimental observation of enhanced activity specifically in the intact hemisphere. This resulted in a persistent left hemispheric bias across frequencies, with a particularly pronounced effect at 9 Hz (Figure 9B). In the third configuration, we set different natural frequencies by lowering the right hemisphere (ω_R_ = 7 Hz). This configuration showed a significant right hemispheric bias at 6 Hz, close to the natural frequency of the right hemisphere, and a left hemispheric bias at 9 Hz (Figure 9C). Notably, when we removed the external periodic force in the first configuration, no hemispheric bias was observed (see red bar in Figure 9A), indicating that the asymmetric coupling manifests its effects only through interaction with external stimulation.

**Figure 9.**
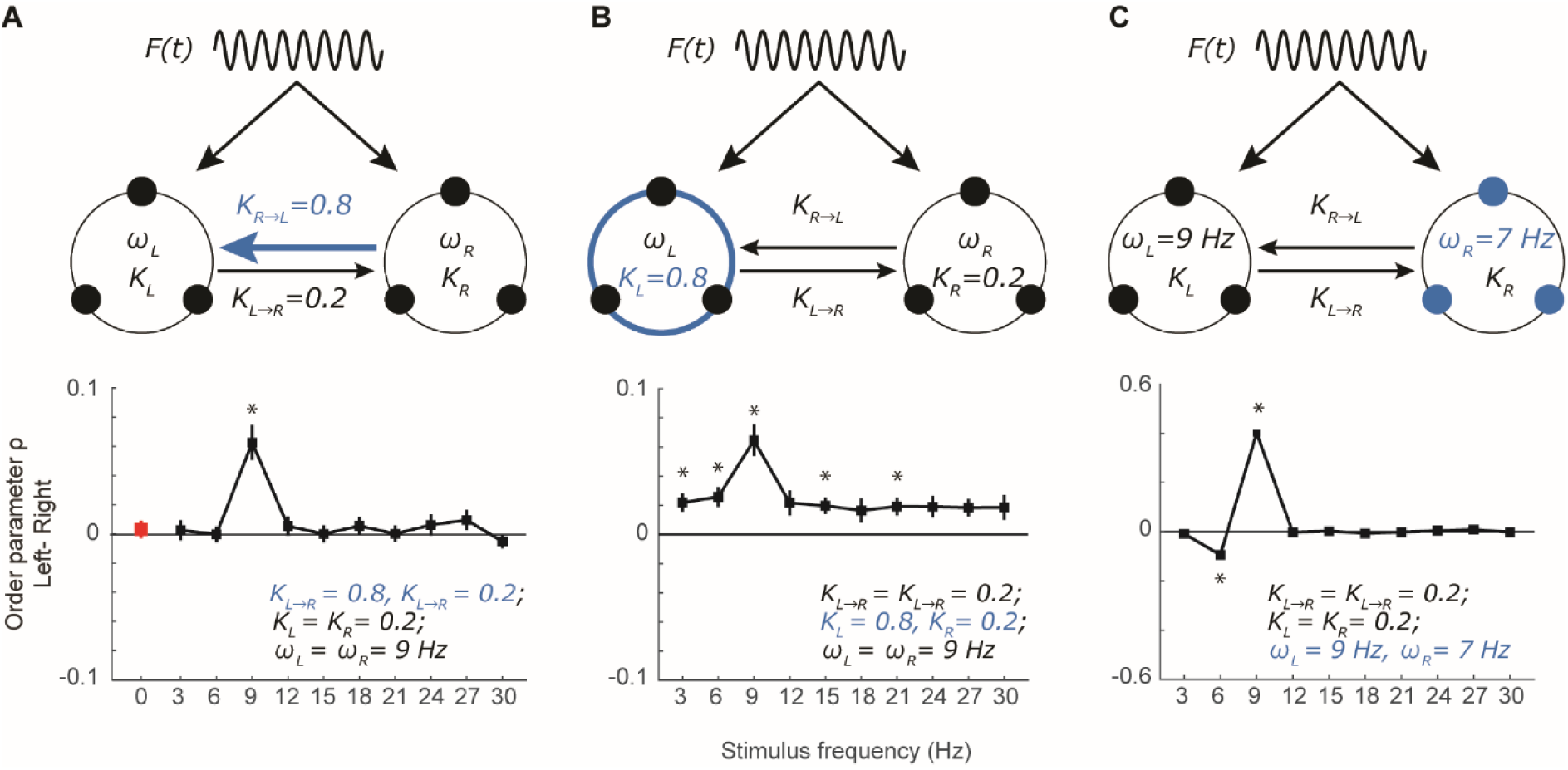
Hemispheric imbalance in model simulations. Upper panels **(A-C)** show schematic illustrations of three different model configurations. Lower panels show corresponding order parameter differences between hemispheres across stimulus frequencies. **(A)** Model with asymmetric interhemispheric coupling (*K_R→L_* = 0.8, *K_L→R_* = 0.2) reproduced the experimental findings, showing significant hemispheric bias at 9 Hz. The red bar indicates the values without external stimulation (*F(t)* = 0). **(B)** Model with enhanced left intrahemispheric coupling (*K_L_* = 0.8, *K_R_* = 0.2) produced a persistent bias across frequencies. **(C)** Model with different natural frequencies between hemispheres (*ω_L_* = 9, *ω_R_* = 7) led to frequency-specific biases at both hemispheres’ natural frequencies. Asterisks indicate significant differences (\**p* < 0.05, FDR corrected). Error bars represent 2 standard errors

## Discussion

Our experiments revealed that while SSVEPs can be reliably elicited by flickering stimuli at various frequencies, only the USN group exhibited a significant hemispheric difference in SSVEP at 9 Hz. The computational model demonstrated that this hemispheric bias could arise from both the resonance between intrinsic and stimulus frequencies and stronger right-to-left interhemispheric coupling. In contrast to the healthy control group, this difference was attributed to an increase in activity within the intact hemisphere rather than a decrease in the affected hemisphere. The enhanced response in the intact hemisphere was accompanied by increased phase-amplitude coupling between alpha and gamma oscillations, suggesting that the entrained alpha oscillations systematically modulated local high-frequency neural activity.

Furthermore, the increased information flow from the right visual cortex to the left frontal cortex revealed by TE analysis is consistent with the model predictions, and these results indicate an imbalance in information flow between hemispheres in addition to changes in local activity.

### Alpha frequency-specific hemispheric bias and entrainment in USN patients

We observed hemispheric bias in stimulus-locked responses to flickering stimuli in USN patients in the alpha band, which corresponds to the natural frequency of the visual system (Rosanova et al. 2009; Okazaki et al. 2021). The entrainment of the oscillatory system to an external stimulus is caused by the periodic phase shift of neural oscillations to the delivered pulse (Lakatos, Gross, and Thut 2019). At the EEG level, this is expressed as an increase in amplitude at the stimulus frequency as multiple oscillators begin to cycle in synchrony (Thut, Schyns, and Gross 2011).

In general, entrainment is particularly effective when the stimulation period closely matches the oscillator’s natural frequency (Pikovsky, Rosenblum, and Kurths 2003). This principle also applies to neural oscillations: rhythmic stimulation at frequencies that resonate with physiological rhythms influences cognitive and neural function by synchronizing task-related oscillations (Klimesch, Sauseng, and Gerloff 2003; Sauseng et al. 2009; Romei et al. 2011). Our previous research demonstrated that entrainment is most effective when the stimulation frequency closely matches the individual alpha frequency (Okazaki et al. 2021). We also observed that different brain regions have distinct natural frequencies, with entrainment being most pronounced in the alpha band in the visual cortex. Therefore, the observed bias in the 9-Hz frequency-specific SSVEP in USN patients does not merely reflect a hemispheric difference in evoked visual responses to rhythmic external stimuli but, rather, a bias in the synchronous function of intrinsic neural oscillators in the visual area at a specific frequency band (i.e., the alpha band).

Our computational model provides a theoretical framework for understanding these hemispheric differences. Among the three potential mechanisms explored, the model with asymmetric interhemispheric coupling (*K_R→L_ > K_L→R_*) best reproduced the frequency-specific hemispheric bias observed in our experiments. While the coupling parameter *K* in our phase oscillator model simply determines the strength of phase synchronization between hemispheres, this asymmetry might reflect the underlying imbalance in interhemispheric inhibition. Indeed, paired-pulse TMS studies have directly demonstrated enhanced inhibitory influence from the intact to the affected hemisphere in stroke patients (Murase et al. 2004). This interhemispheric imbalance has been further supported by evidence of altered functional connectivity and hyperexcitability in the intact hemisphere (Koch et al. 2008; Grefkes and Fink 2011).

In this model, both hemispheres with a natural frequency of 9 Hz resonate with stimulation at the same frequency, but due to the stronger right-to-left coupling, the resonant response in the right hemisphere more strongly drives oscillations in the left hemisphere. This suggests that the hemispheric bias observed in USN patients is driven by asymmetric interhemispheric coupling of neural networks underlying alpha oscillations. This model prediction also aligns with our experimental observation of enhanced transfer entropy from the right visual cortex to the left frontal areas in USN patients.

Furthermore, it demonstrates that while the system maintains balanced activity between hemispheres in the absence of external stimulation, the underlying asymmetry becomes apparent with external input. This explains why spontaneous alpha oscillations remain normal despite the emergence of a specific bias in response to 9 Hz stimulation, providing a theoretical basis for why USN symptoms become more pronounced during sensory processing and attentional demands.

### Local hemispheric bias of alpha-band entrainment and attentional dysfunction in USN patients

Hyperactivity of the SSVEP and PAC in the left visual cortex in stroke patients may result from a lack of hemispheric reciprocal inhibition (Murase et al. 2004). This scenario is consistent with the concept that in USN, decreased inhibition from the affected right hemisphere causes overactivity of the visual cortex within the intact left hemisphere, which results in a left hemisphere bias in visual processing (Corbetta and Shulman 2011; Koch et al. 2008). However, the hemispheric bias observed in the present study was frequency-specific and was not observed in non-USN patients (who also had right hemisphere damage), suggesting that it may reflect not only the overactivity of the visual cortex but also alterations in the neural networks’ capacity to synchronize with external stimuli at specific frequencies. The entrainment of neural oscillations to external stimuli is achieved via consistent phase alignment, which, in turn, systematically influences the timing of the higher frequency components of neural activity (Glim et al. 2019). Consistent with this, we observed enhanced PAC between alpha phase and gamma amplitude in the intact hemisphere of USN patients, supporting the interpretation that entrained alpha oscillations systematically modulate high-frequency activity.

This phase coordination is influenced by both external factors and internal processes, such as attention in anticipatory contexts. Attention adjusts the temporal sequence of excitatory and inhibitory phases in neural oscillations, thereby influencing perception (Lakatos et al. 2008; Schroeder and Lakatos 2009; Besle et al. 2011; Bonnefond and Jensen 2012). Given the importance of alpha-band oscillations in attentional processes (Klimesch 2012), the hemispheric bias in oscillatory synchrony observed in USN patients may reflect critical abnormalities in cortical excitability coordination. Therefore, the hemispheric bias in synchronization capacity at frequencies relevant to attention likely disrupts the coordinated oscillatory activity necessary for balanced spatial attention processing.

Clinically, USN presents as neglect of the left visual field and is generally thought to reflect a relative dominance of rightward orienting. Because alpha power is often interpreted as reflecting functional inhibition (Worden et al. 2000; Kelly et al. 2006; Foxe and Snyder 2011), classic alpha-power lateralization associated with rightward orienting would typically predict increased alpha power over the right hemisphere and decreased alpha power over the left. However, in our data, right-hemisphere alpha power during stimulation was comparable to that of controls, whereas the intact left hemisphere exhibited stronger stimulus-locked synchronization (SSVEP) and enhanced alpha–gamma coupling. Thus, the present findings do not appear to reflect a simple amplification of tonic spatial alpha-power lateralization. Rather, they suggest a dynamic, temporally selective form of alpha-mediated inhibition that organizes the timing of local excitability. From this perspective, alpha-power lateralization and stimulus-locked entrainment may reflect related but distinct aspects of alpha-based inhibitory control, with the former regulating spatial gating and the latter regulating the timing of sensory processing (Jensen and Mazaheri 2010).

Moreover, such a timing-based bias may arise even in the absence of explicit flicker. Visual input is continuously sampled under the influence of intrinsic alpha activity, and perceptual sensitivity fluctuates with the phase of ongoing oscillations (Romei et al. 2008; Iemi et al. 2017; VanRullen 2016). As a consequence, processing is relatively facilitated when incoming events coincide with high-excitability phases and relatively suppressed otherwise (Busch, Dubois, and VanRullen 2009; Mathewson et al. 2010). Thus, a hemispheric bias in alpha-band synchronization capacity could bias the timing with which sensory input is sampled across the two hemispheres, even without explicit rhythmic stimulation. This view is also consistent with the possibility that the bias is less apparent during eyes-closed rest with minimal visual input, yet becomes behaviorally expressed in everyday settings where continuous input is sampled in a phase-dependent manner (Landau and Fries 2012; VanRullen 2016). Overall, a hemispheric bias in synchronization capacity within the alpha range likely disrupts the temporally coordinated sampling of visual input required for balanced spatial attention, contributing to the attentional deficits observed in USN.

### Biased information transfer in USN patients

Locally, the USN group showed a higher SSVEP and greater PAC in the left visual cortex, whereas TE from the right visual cortex to the left frontal cortex was higher than TE from the left visual cortex to the left frontal cortex. This hemispheric difference appeared only in the Feedforward (visual-to-frontal) direction, and no such difference was observed in the Feedback (frontal-to-visual) direction. This directional asymmetry argues against spurious overestimation of TE due to autocorrelation or volume conduction (Daube, Gross, and Ince 2022), because such pseudo-causal effects would be expected to manifest more symmetrically in both directions. In addition, by computing TE from the amplitude envelope of the 9-Hz activity, we reduced the influence of strong autocorrelation inherent in narrow-band oscillatory signals and thereby minimized the conditions that Daube et al. identified as leading to TE overestimation. Taken together, these points suggest that the observed TE asymmetry is unlikely to be explained solely by methodological artifacts and may reflect a genuine directional imbalance in interregional communication following right-hemisphere damage.

Given that right frontal damage is one of the common causes of USN, this difference may be an adaptive neural response to maintain hemispheric communication and compensate for attentional dysfunction. Specifically, input from the right visual cortex may bypass the affected right frontal cortex to enable enhanced information flow to the intact left frontal cortex, thereby supporting attentional functions. This compensatory strategy of increased information flow is consistent with functional imaging study findings on the reorganization of functional networks following stroke; moreover, it supports the idea of neuroplasticity, whereby patients can recover functions and maintain cognitive abilities following neurological injury (Rehme et al. 2011; Nudo 2006; Saur et al. 2006). This interpretation is also consistent with the trend-level correlation shown in Supplementary Figure 3C. Although the correlation did not survive correction for multiple comparisons, patients with a stronger feedforward TE bias toward the left frontal region tended to show higher BIT scores. This observation raises the possibility that asymmetric information transfer from the right visual cortex to the intact left frontal cortex may contribute to compensatory network reorganization associated with milder neglect symptoms.

### Distinct neural responses in non-USN and USN patients

Although both the non-USN and USN groups had right hemisphere damage, they exhibited distinct neural responses to alpha-band rhythmic stimulation. In non-USN patients, the observed bilateral enhancement of alpha-band synchrony likely reflects a compensatory recruitment of residual right-hemisphere circuits that are still able to cooperate with the intact left hemisphere. Neuroimaging studies have reported that patients who show this kind of bilateral reorganization pattern early after stroke tend to achieve better recovery outcomes (Ward et al. 2003; Rehme et al. 2011; Grefkes and Fink 2011). Thus, the bilateral pattern seen in the non-USN group may correspond to such a favorable reorganization profile, and we propose that engaging both hemispheres in parallel helps maintain frequency-specific rhythm coordination and protects against the development of spatial neglect. In contrast, the neural responses in USN patients are more asymmetric, with a local increase in alpha-band SSVEP primarily observed in the intact left visual cortex. This asymmetry may result from the damaged right hemisphere being less effectively engaged, leading to an over-reliance on the left hemisphere (Corbetta et al. 2005). This suggests that the right hemisphere damage in USN patients might affect more critical parts of the attentional network.

To further characterize the anatomical background of these group differences, we generated lesion overlap maps and quantified lesion volume in the USN and non-USN groups (Supplementary Figure 1). Both groups predominantly showed lesions involving the right MCA territory, but lesion volume was significantly larger in the USN group than in the non-USN group (USN: 72,419 ± 73,778 mm³; non-USN: 11,739 ± 23,907 mm³; Mann–Whitney *U* = 361.0, *p* = 1.41 × 10⁻⁵). These anatomical differences indicate that lesion extent is an important factor associated with USN. However, they do not by themselves fully explain the frequency-specific neural effect observed here. Importantly, this frequency specificity coincides with the intrinsic alpha frequency of the visual system. This correspondence suggests that the present finding may not simply arise from lesion location or lesion volume alone, but may instead reflect a more complex mechanism involving selective functional disruption of oscillatory networks. From this perspective, lesion topography and oscillatory dynamics should be regarded not as competing explanations, but as different levels at which the same pathological condition can be understood. The key question is which network components are affected and how their dysfunction gives rise to the frequency-selective hemispheric imbalance in synchronization capacity at 9 Hz. Thus, even if USN and non-USN patients differ in lesion extent and aspects of lesion topography, this does not undermine the present interpretation, but rather highlights the need to examine how anatomical damage relates to frequency-specific network dysfunction.

Our computational model illustrates a plausible mechanism for such a process. When two coupled oscillators sharing the same intrinsic alpha frequency are connected via asymmetric interhemispheric couplings, the model selectively produces an imbalance in synchrony at the resonant frequency, whereas responses at non-resonant stimulation frequencies remain relatively balanced between hemispheres. This model result suggests that post-lesion asymmetry in interhemispheric coupling may bias alpha-band information processing (e.g. phase-dependent sampling/synchrony) between hemispheres, and may consequently manifest as systematic biases in perceptual and attentional allocation.

Interestingly, although right-hemisphere alpha-band SSVEP power did not differ significantly between the USN group and healthy controls, it showed a significant positive correlation with BIT scores within the USN group. This result indicates that, even when the quantitative power of right-hemisphere alpha oscillations appears normal, their qualitative contribution to the spatial-attention network is altered in USN. Such a qualitative dissociation, together with compensatory hyperactivity in the intact left hemisphere and a rerouting of information flow from the right visual cortex to the left frontal cortex, likely skews the overall balance of spatial attention. The interplay among these quantitative factors, and between these dynamics and their behavioral consequences, is probably nonlinear, and the present data cannot fully unravel their complex interactions; nevertheless, the principle that “normal power ≠ normal function” strongly suggests that USN arises from a qualitative breakdown in rhythm coordination, offering a new perspective on the neural basis of unilateral spatial neglect.

### Limitations

This study has several limitations that must be acknowledged. First, although we revealed a 9-Hz-specific hemispheric bias in USN, the study does not pinpoint where within the cortical network the abnormal synchrony originates. Higher spatial-resolution techniques, such as magnetoencephalography or high-density intracranial recordings, could map the generators and propagation pathways of alpha-band entrainment more precisely and thus identify the circuit nodes in which synchrony breaks down. Second, although we added lesion overlap maps and quantified lesion volume, these analyses primarily characterize where and how extensively lesions were distributed across the two patient groups. They do not directly identify which anatomical network disruptions give rise to the 9 Hz-specific imbalance in interhemispheric synchrony. Future studies with larger cohorts will be necessary to combine detailed lesion-symptom mapping and assessments of white-matter disconnection with neural synchrony analyses to determine which anatomical network disruptions lead to frequency-specific alterations in oscillatory dynamics after stroke. Third, the cross-sectional design precludes definitive causal inferences. Longitudinal studies that track changes in alpha-band asymmetry and behavioral recovery over time, or before and after rehabilitation interventions, will be essential to determine how dynamic reorganization of neural synchrony relates to the evolution and remediation of spatial neglect. Fourth, the hemispheric bias in alpha–gamma PAC should be interpreted in light of potential contamination from miniature saccades (Yuval-Greenberg et al. 2008). However, several observations make it unlikely that such artifacts are the primary source of the effect. For saccadic spike potentials to account for the PAC under 9-Hz stimulation, they would need to occur in a highly periodic and phase-locked manner relative to the 9-Hz cycle. This scenario is unlikely given the stimulus-locked dynamics of miniature saccades (i.e., post-stimulus inhibition followed by a rebound around 200–300 ms) (Yuval-Greenberg and Deouell 2009). Moreover, SP-related contamination would be expected to yield a broadband gamma profile (∼20–90 Hz) (Keren, Yuval-Greenberg, and Deouell 2010; Yuval-Greenberg and Deouell 2009), rather than the relatively narrow band (35–45 Hz) observed here. In addition, we found that gamma-band power in the 35–45 Hz range did not show any hemispheric difference between O1 and O2 during 9-Hz stimulation (data not shown).

## Conclusions

Our study demonstrates fundamental differences in neural responses between stroke patients with USN and without USN. Empirical data and computational modeling together show that asymmetric interhemispheric coupling, resonating with the brain’s intrinsic alpha rhythm, selectively amplifies 9-Hz entrainment in the intact hemisphere of USN patients. This amplification is accompanied by stronger α–γ phase–amplitude coupling and asymmetric information flow, indicating that the effect propagates beyond local oscillations to reconfigure cross-frequency interactions and large-scale network dynamics. Thus, USN can be regarded as a disorder in which a qualitative breakdown of alpha-band rhythm coordination destabilizes the dynamic equilibrium of the attention network. Longitudinal studies that track the plastic evolution of these synchrony patterns during recovery and refine frequency-specific neuromodulation strategies will be crucial for translating these insights into therapy.

## Acknowledgments

This work was supported in part by the NINS OPEN MIX LAB Program (NINS Program No. OML032401); a research grant from TOYOTA Motor Corporation to KK; and a grant-in-aid for General Research on Adult Diseases Including Cardiovascular Diseases and Diabetes (23FA1015) from the Japanese Ministry of Health, Labor and Welfare to IM. We also thank Dr. Hiroshi Yokoyama for providing the code for the transfer entropy analysis.

## Data, Materials, and Software Availability

Upon reasonable request, the anonymized data may be made available subject to institutional approval, and analysis code and simulation scripts used in this study are available from the corresponding author

## Supplementary figure

**Supplementary Figure 1:**
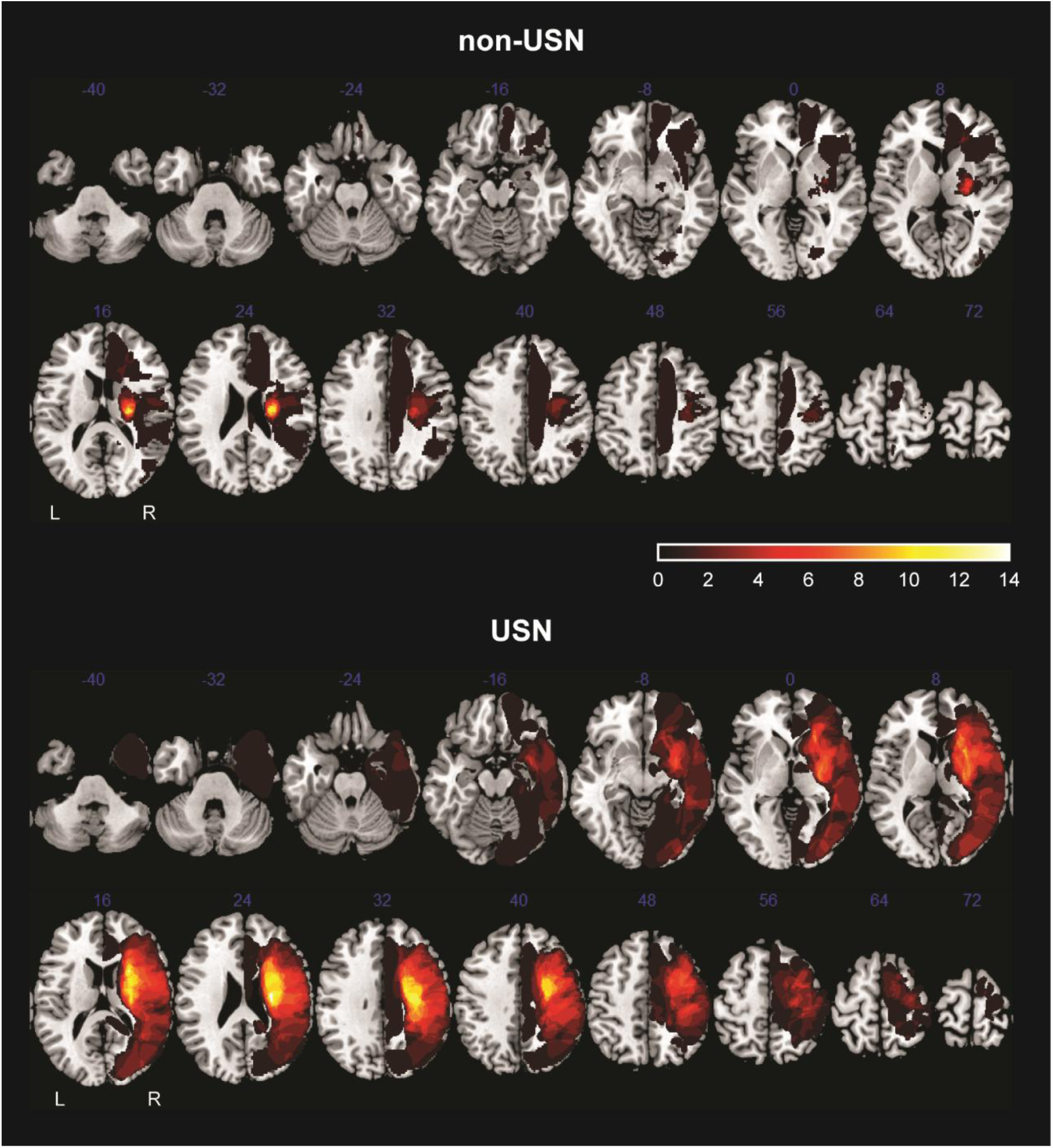
Lesion overlap maps in non-USN and USN patients. Maps are shown for non-USN patients (top) and USN patients (bottom), with warmer colors indicating a greater number of overlapping lesions at each voxel. MRI was performed on a 1.5-T scanner, and T2-weighted and 3D T1-weighted whole-brain images were acquired. Stroke lesions were manually delineated on T2-weighted MRI and normalized to Montreal Neurological Institute (MNI) space using the Clinical Toolbox (Rorden et al. 2012) in SPM8 (Tierney et al. 2025). Acquisition details are reported elsewhere (Kawano et al. 2020). Both groups predominantly showed lesions involving the right MCA territory, but lesion volume was significantly larger in the USN group than in the non-USN group (USN: 72,419 ± 73,778 mm³; non-USN: 11,739 ± 23,907 mm³; Mann–Whitney U = 361.0, p = 1.41 × 10⁻⁵).

**Supplementary Figure 2:**
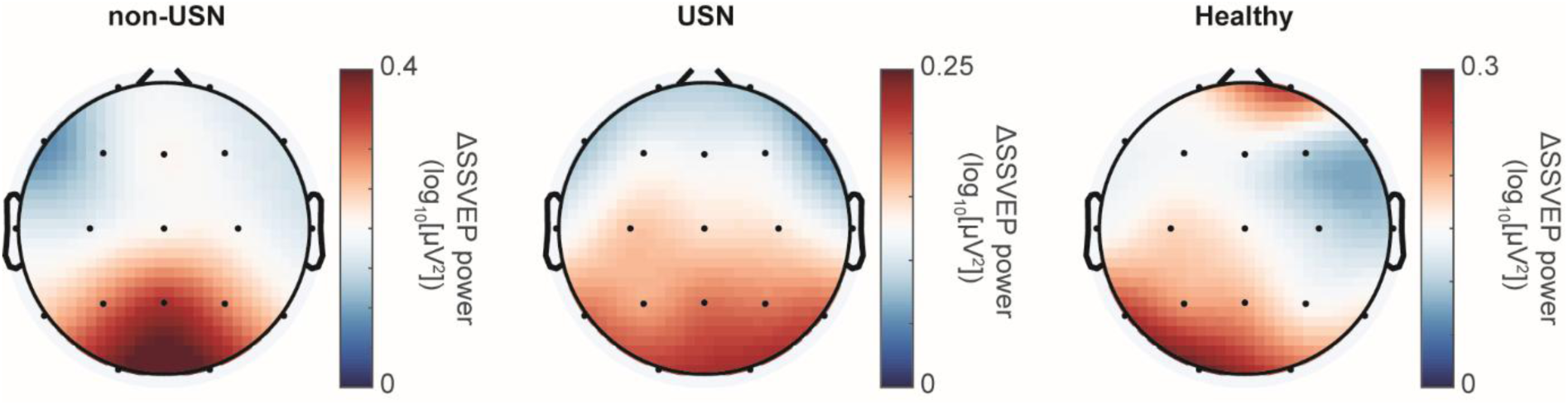
Scalp distribution of SSVEP power across groups. Topographical maps show EEG power at the fundamental (stimulation) frequency in the non-USN, USN, and healthy control groups. Power at each frequency was log10-transformed, baseline-corrected by subtracting the pre-stimulus period (−9.5 to −0.5 s), and then averaged across stimulation frequencies and participants. In all groups, the strongest responses were observed over occipital electrodes (O1 and O2).

**Supplementary Figure 3:**
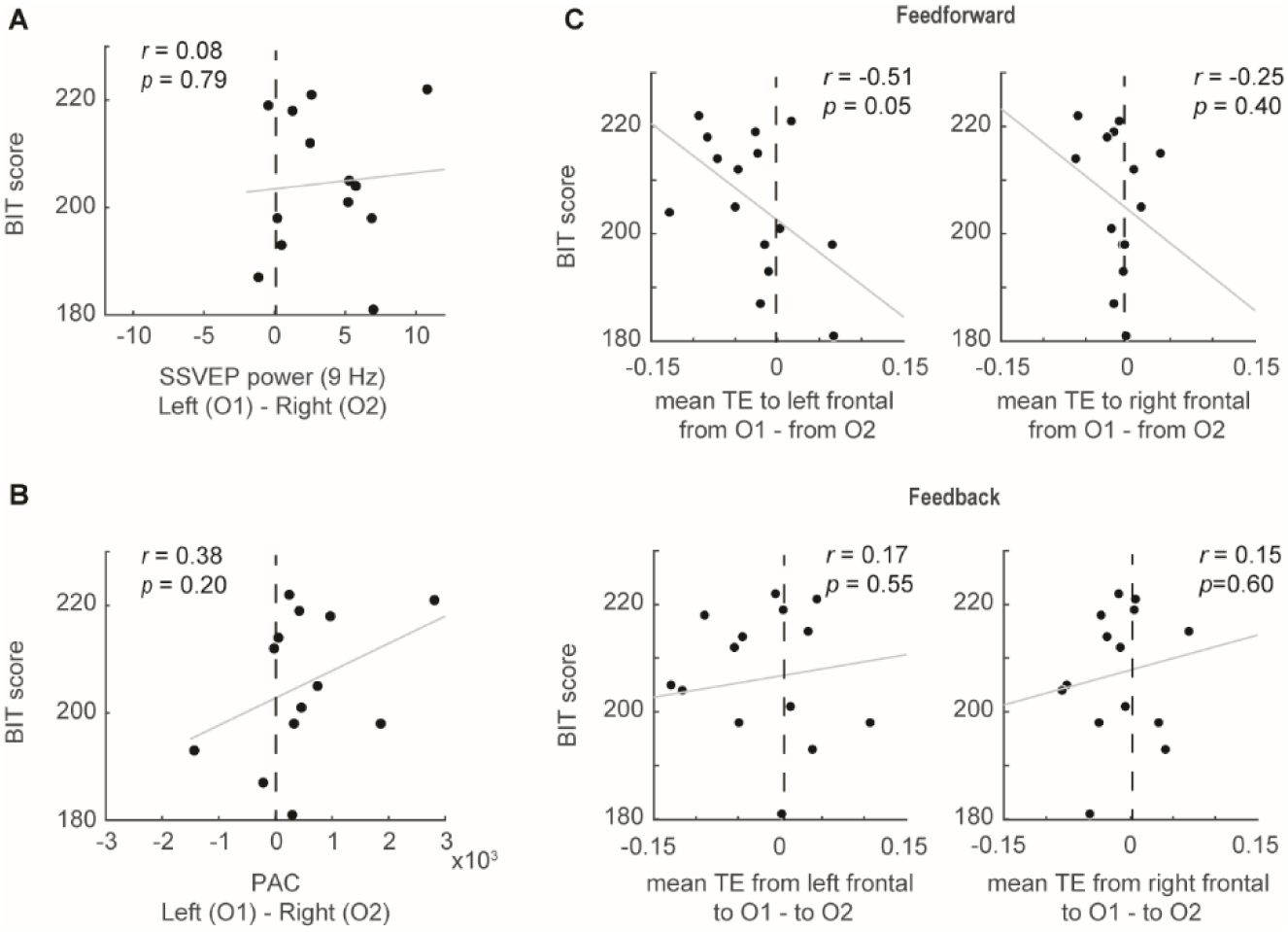
Correlations between BIT score and hemispheric imbalance measures during 9-Hz stimulation in USN patients. (A) Correlation between BIT score and the hemispheric imbalance of 9-Hz SSVEP power shown in Figure 3. (B) Correlation between BIT score and the hemispheric imbalance of PAC. PAC was calculated as the modulation index between the 9-Hz phase and gamma-band amplitude (33–46 Hz), corresponding to the significant cluster shown in Figure 6, and averaged across this gamma-frequency range. (C) Correlation between BIT score and the hemispheric imbalance of TE shown in Figure 7, separately for feedforward and feedback interactions.

